# Medium Composition Affects Microbial Corrosion Rates

**DOI:** 10.1101/2024.04.11.589044

**Authors:** Di Wang, Toshiyuki Ueki, Peiyu Ma, Dake Xu, Derek R. Lovley

## Abstract

*Desulfovibrio vulgaris* and *Desulfovibrio ferrophilus* were previously proposed to have distinct iron corrosion mechanisms because *D. ferrophilus* corroded faster. However, the chloride concentration in the *D. ferrophilus* ‘marine’ medium was much higher than in the *D. vulgaris* ‘freshwater’ medium. *D. vulgaris* corrosion rates accelerated with increasing chloride and were faster than *D. ferrophilus* in the same marine medium. Differences in *D. ferrophilus* corrosion rates in two different media with the same chloride concentration suggested that minor differences in other medium constituents also impact on microbial corrosion. These results demonstrate the importance of considering medium composition in microbial corrosion studies.

## Introduction

Microbial activity promoting the corrosion of iron-containing metals has an important economic and safety impact on a wide range of infrastructure, water treatment and distribution systems, gas and oil pipelines, and medical devices [1–3]. Sulfate-reducing bacteria are major contributors to metal corrosion in diverse anaerobic environments and have been the primary model microbes in corrosion mechanistic studies [1, 3–6]. *Desulfovibrio vulgaris* has been the primary pure culture model microbe for the development of many of the concepts for how sulfate reducers promote corrosion [6–13]

Studies with a hydrogenase-deficient mutant of *D. vulgaris* unable to utilize H_2_ have demonstrated that H_2_ serves as an intermediary electron transfer to facilitate electron transfer between Fe^0^ and cells [8, 14]. Even in the absence of microbial cells Fe^0^ anaerobically corrodes as the result of Fe^0^ oxidation coupled to the reduction of H^+^ to H_2_:

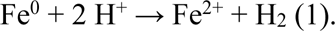

*D. vulgaris* oxidizes H_2_ with the reduction of sulfate, generating sulfide:

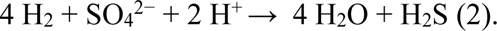

The generation of sulfide further stimulates H_2_ production. For example, sulfide combines with Fe^2+^ to form ferrous sulfide that deposits on metal surfaces and promotes the electron transfer in reaction #1 [3]. Furthermore, hydrogen sulfide can react with Fe^0^ to generate H_2_ [1, 4]:

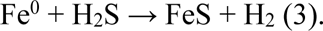

Thus sulfide-facilitated H_2_ production from reactions #1 and #3 functions as a positive feedback loop leading to even more sulfide generation from reaction #2, which in turn produces more H_2_.

Some microbes are capable of obtaining electrons from Fe^0^ via direct Fe^0^-to-microbe electron transfer (i.e. electrobiocorrosion [1]). Electrobiocorrosion has been rigorously demonstrated in several *Geobacter* species [15, 16], *Shewanella oneidensis* [17, 18], and *Methanosarcina acetivorans* [19]. Strains in which genes for key outer-surface *c*-type cytochromes were deleted had diminished rates of Fe^0^ oxidation and strains unable to utilize H_2_ continued to promote Fe^0^ oxidation. Several electrobiocorrosion studies demonstrated that 316L stainless steel was a good Fe^0^ source to evaluate whether microbes are capable of direct electron uptake because, unlike pure Fe^0^, it does not generate H_2_ to support the growth of microbes that rely on H_2_ as an intermediary electron carrier [16, 19]. Investigations with stainless steel have identified additional *Methanosarcina*, as well as *Methanothrix*, and *Sporomusa* species that also appear to be capable of electrobiocorrosion [20].

Current evidence suggests that *D. vulgaris* is not capable of electrobiocorrosion. Stainless steel did not support *D. vulgaris* sulfate reduction [8]. The *D. vulgaris* hydrogenase-deficient mutant did not reduce sulfate with Fe^0^ as the sole electron donor, further demonstrating the need for H_2_ to serve as an intermediary electron carrier between Fe^0^ and cells [8]. Furthermore, there was 1:1 correspondence between Fe^0^ loss and H_2_ accumulation in the mutant cultures, consistent with Fe^0^ oxidation coupled to H_2_ production (reaction #1) [14]. If electrobiocorrosion was active, then the loss of Fe^0^ would have been greater than H_2_ production.

Fe^0^ was corroded faster in the presence of *Desulfovibrio ferrophilus* than the closely related *D. vulgaris* [21]. It was inferred that the higher corrosion rates could be attributed to *D. ferrophilus* electrobiocorrosion capabilities not found in *D. vulgaris* [4, 21, 22]. However, there is no experimental evidence that electrobiocorrosion is a faster mechanism for Fe^0^ oxidation than H_2_ production (Xu et al. 2023). In fact, *D. ferrophilus* was unable to reduce sulfate or other electron acceptors with stainless steel, a suitable electron donor for all microbes rigorously shown to be capable of electrobiocorrosion [23]. Fe^0^ was a suitable electron donor for *D. ferrophilus*, suggesting that like *D. vulgaris*, it requires H_2_ as an intermediary electron carrier [23]. Other microbes that were isolated along with *D. ferrophilus* and proposed to directly extract electrons from Fe^0^ have also been subsequently found to depend on H_2_ as an electron shuttle [20, 23]. These results and other studies [24] have demonstrated that the previous assumption [4, 21, 22] that fast rates of Fe^0^ corrosion are indicative of a direct electron uptake mechanism is not warranted.

The evidence that, like *D. vulgaris*, *D. ferrophilus* obtains electrons from Fe^0^ via H_2_ suggests that factors other than a different mechanism for electron uptake from Fe^0^ account for the faster corrosion of iron-containing metals in the presence of *D. ferrophilus*. A significant difference between *D. ferrophilus* and *D. vulgaris* studies was the concentration of chloride in the media. *D. ferrophilus* was grown in ‘marine media’ in which the addition of sodium chloride and magnesium chloride provided 473 mM-495 mM chloride [22, 25, 26]. In contrast the chloride concentration in the ‘freshwater’ *D. vulgaris* media was only 12 mM-19 mM [8, 27, 28].

Differences in medium chloride concentrations are relevant because chloride ion greatly accelerates abiotic iron corrosion [29, 30]. Chloride penetrates protective mineral layers and influences corrosion reactions and corrosion products to favor faster overall corrosion and/or greater pitting [31–33]. The impact of chloride concentrations on microbial corrosion was previously studied with the halophilic archaeon *Natronorubrum tibetense*, but the primary stimulatory effect of chloride appeared to be this halophile’s requirement for high chloride concentrations for growth [34]. Increasing the chloride concentration further inhibited corrosion because chloride interfered with *N. tibetense* attachment to metal and abiotic corrosion reactions [34]. These results demonstrate the potential complexities in evaluating the role of chloride in microbial corrosion.

We hypothesized that differences in rates of Fe^0^ corrosion between *D. ferrophilus* and *D. vulgaris*, might simply be due to differences in media chloride concentrations. *D. vulgaris* has previously been adapted to grow at chloride substantially higher than those in typical *D. vulgaris* culture medium [35]. Our results demonstrate that *D. vulgaris* corroded Fe^0^ faster in high chloride medium than in low chloride medium. *D. ferrophilus* corrosion rates were lower than *D. vulgaris* corrosion rates when the two were incubated in the same medium. Thus, medium composition is a key consideration when comparing corrosion rates between microbes or between laboratory cultures and actual corrosion sites.

## Materials and Methods

### Cultures and growth medium

*D. vulgaris* strain JW710 was obtained from a repository of *D. vulgaris* strains maintained by Valentine V. Trotter and Adam M. Deutschbauer of the Lawrence Berkeley Laboratory, Berkeley, CA USA. *D. ferrophilus* was obtained from DSMZ-German Collection of Microorganisms and Cell Cultures GmbH. Cultivation followed standard anaerobic procedures. Oxygen was removed from all gas streams, open bottles were maintained under a stream of gas, or in a Coy anaerobic glove bag (Coy Laboratory Products Inc., MI, USA). Culture vessels were sealed with thick butyl rubber stoppers. Gas-flushed needles and syringes were employed for all culture sampling and transfers. Incubations were at 30 °C.

*D. vulgaris* was grown as previously described [8] in NB medium which contains (per liter of deionized water): 0.42 g of KH_2_PO_4_, 0.22 g of K_2_HPO_4_, 0.2 g of NH_4_Cl, 0.38 g of KCl, 0.36 g of NaCl, 0.04 g of CaCl_2_·2H_2_O, 0.1 g of MgSO_4_·7H_2_O, 1.8 g of NaHCO_3_, 0.5 g of Na_2_CO_3_, 1.0 mL of 1 mM Na_2_SeO_4_, 15.0 mL of a vitamin solution [20], and 10.0 mL of NB trace mineral solution. The composition of the NB trace mineral solution per L of deionized water is 2.14 g of nitrilotriacetic acid, 0.1 g of MnCl_2_·4H_2_O, 0.3 g of FeSO_4_·7H_2_O, 0.17 g of CoCl_2_·6H_2_O, 0.2 g of ZnSO_4_·7H_2_O, 0.03 g of CuCl_2_·2H_2_O, 0.005 g of AlK(SO_4_)_2_·12H_2_O, 0.005 g of H_3_BO_3_, 0.09 g of Na_2_MoO_4_, 0.11 g of NiSO_4_·6H_2_O, and 0.02 g of Na_2_WO_4_·2H_2_O. Cultures were routinely grown with 60 mM DL lactate as electron donor and 30 mM sulfate as the electron acceptor. Additional sodium chloride was added to increase the chloride concentration as indicated in the Results section.

*D. ferrophilus* was grown in either NB medium in which the chloride concentration was increased to 400 mM or in 195c medium [23]. The 195c medium contains (per liter of deionized water): 4.26 g of Na_2_SO_4_, 21 g of NaCl, 3.0 g of MgCl_2_·6H_2_O, 10 mL of salt mix including (20 g of KH_2_PO_4_, 30 g of NH_4_Cl and 50 g of KCl per liter), 0.15 g of CaCl_2_, 1.5 g Na_2_CO_3_, 60 mM Sodium DL-Lactate, 10 mL trace mineral solution per L of deionized water is 10 mL of HCl (25%), 1.5 g of FeCl_2_·4H_2_O, 70 mg of ZnC_l2_, 100 mg of MnCl_2_·4H_2_O, 6 mg of H_3_BO_3_, 190 mg of CoCl_2_·6H_2_O, 2 mg of CuCl_2_·2H_2_O, 24 mg of NiCl_2_·6H_2_O, 36 mg of Na_2_MoO_4_·2H_2_O, 10.0 mL of a vitamin solution and 1 mL SeW solution containing 0.50 g NaOH, 3 mg Na_2_SeO_3_·5H_2_O and 4 mg Na_2_WO_4_·2H_2_O per liter. The chloride concentration of this medium is 400 mM.

Growth with lactate as the sole electron donor was evaluated in 10 mL of medium in 28 mL anaerobic pressure tubes under the atmosphere of N_2_/CO_2_ (80:20). All the medium in pressure tubes were bubbled with N_2_/CO_2_ (80:20) for 15 min before the tubes were sealed with a thick butyl rubber stopper. Turbidity in the tubes was measured at 600 nm in a spectrophotometer (Model Genesys 50, Thermo Scientific, USA) to monitor growth.

Iron coupons (8 mm × 8 mm × 5 mm; composition (w/w %): Fe 99.95, C 0.01, S 0.008, P 0.01, Si 0.001, Mn 0.001, Ni 0.001, Cr 0.01 and Al 0.006) served as the Fe^0^ source for corrosion studies. The coupons were polished sequentially with 240, 600, and 1000 grit sand paper and then washed with 75% ethanol and MilliQ water. The cubes were sterilized with 75% ethanol and dried in a UV chamber. Three iron coupons were added to 30 mL of NB medium in 65 mL serum bottles containing 5 mM lactate and 5 mM sulfate in the anaerobic glove bag (gas phase, 7:20:73 H_2_/CO_2_/N_2_) and sealed with butyl rubber stoppers. The serum bottles were removed from the glove bag and flushed with N_2_/CO_2_ (80:20) for 10 min. Corrosion studies were initiated with a 5% inoculum of a culture pregrown in NB or 195C medium at the same chloride concentration with lactate as the electron donor and sulfate as the electron acceptor, as described above.

### Sulfate consumption and hydrogen production

For sulfate determinations culture aliquots (0.4 mL) were diluted 20-fold, filtered (0.22 μm; polyvinylidene difluoride) and analyzed with ion chromatography ((Dionex AS-DV, Thermo Scientific, USA). Headspace H_2_ concentrations were measured with a gas chromatograph (Trace 1310, Thermo Scientific, USA) equipped with a thermal conductivity detector.

### Biofilm Imaging

Iron coupons were gently rinsed with phosphate buffer (pH 7.4) and stained with Live/Dead BacLight™ bacterial staining kit (Molecular Probes L-7012, Thermo Fisher Scientific, USA) for 10 min in the dark. Biofilms on the iron surface were observed by a confocal laser scanning microscope using fluorescent channels (Zeiss model LSM 900, Zeiss, Germany).

### Corrosion analysis

For weight loss determination, three iron coupons were removed from four replicate incubations and corrosion products were removed with Clarke’s solution according to ASTM G1-03 protocol[36]. The coupon surfaces (120 µm × 120 µm scanned area for abiotic corrosion studies and 600 µm × 600 µm scanned area for microbial corrosion studies) were scanned in bright field mode with the same confocal laser scanning microscope used for biofilm analysis. The corrosion pit depth and surface morphologies were evaluated using ConfoMap Premium Software provided by Mountains Technology for ZEISS microscopes. Corrosion products were evaluated with X-ray diffraction (9 kW rotating anode X-ray source (λ∼1.54Å) Rigaku SmartLab, Japan). Elemental composition of the coupon surfaces was analyzed with energy-dispersive X-ray spectroscopy (X-Max detector, Oxford Instruments, UK) equipped in a scanning electron microscope (EVO 10, Zeiss, Germany).

### Electrochemical tests

Electrochemical corrosion tests were performed in 500 mL anaerobic glass cells. The 237.5 mL medium and 12.5 mL (5%) culture of each salinity were added in the Coy glove bag. A platinum plate (10 mm × 10 mm × 1 mm) was the counter electrode, a saturated calomel electrode served as the reference electrode, and the pure iron with a 1 cm^2^ square-shaped area exposed, connected with a rubber coated copper wire and sealed with epoxy resin was the working electrode. The glass cell was sealed using a screw lid and all the joints were sealed with paraffin to maintain the anaerobic conditions. All the electrochemical tests of open circuit potential (OCP), linear polarization resistance (LPR), potential dynamic polarization (PDP) measurements were obtained from electrochemical workstations (Reference 600, Gamry Instruments, USA). LPR tests were scanned with a rate of 0.167 mV/s in the range of −10 mV to 10 mV versus OCP. *R*_p_ was calculated as the linear slope of the region within ± 5 mV versus OCP. The PDP curves were collected at a scan rate of 0.333 mV/s from 0 mV to −0.3 V versus OCP and 0 mV to 0.3 V, respectively. The corrosion current density *i*_corr_ was calculated from the extrapolation of the PDP curves.

## Results and Discussion

### Higher Chloride Increases Fe^0^ oxidation coupled to H_2_ production

Increasing the chloride concentration from a ‘freshwater medium’ concentration of 12 mM to a ‘marine medium’ level of 400 mM increased abiotic H_2_ production in sterile medium and the loss of Fe^0^ from the coupons (Fig. 1A, B). This result suggested that there was more Fe^0^ oxidation coupled to H^+^ reduction (reaction #1) in the presence of more chloride. Higher chloride also increased the extent and depth of abiotic pitting (Fig. 1C-I).

**Fig. 1.**
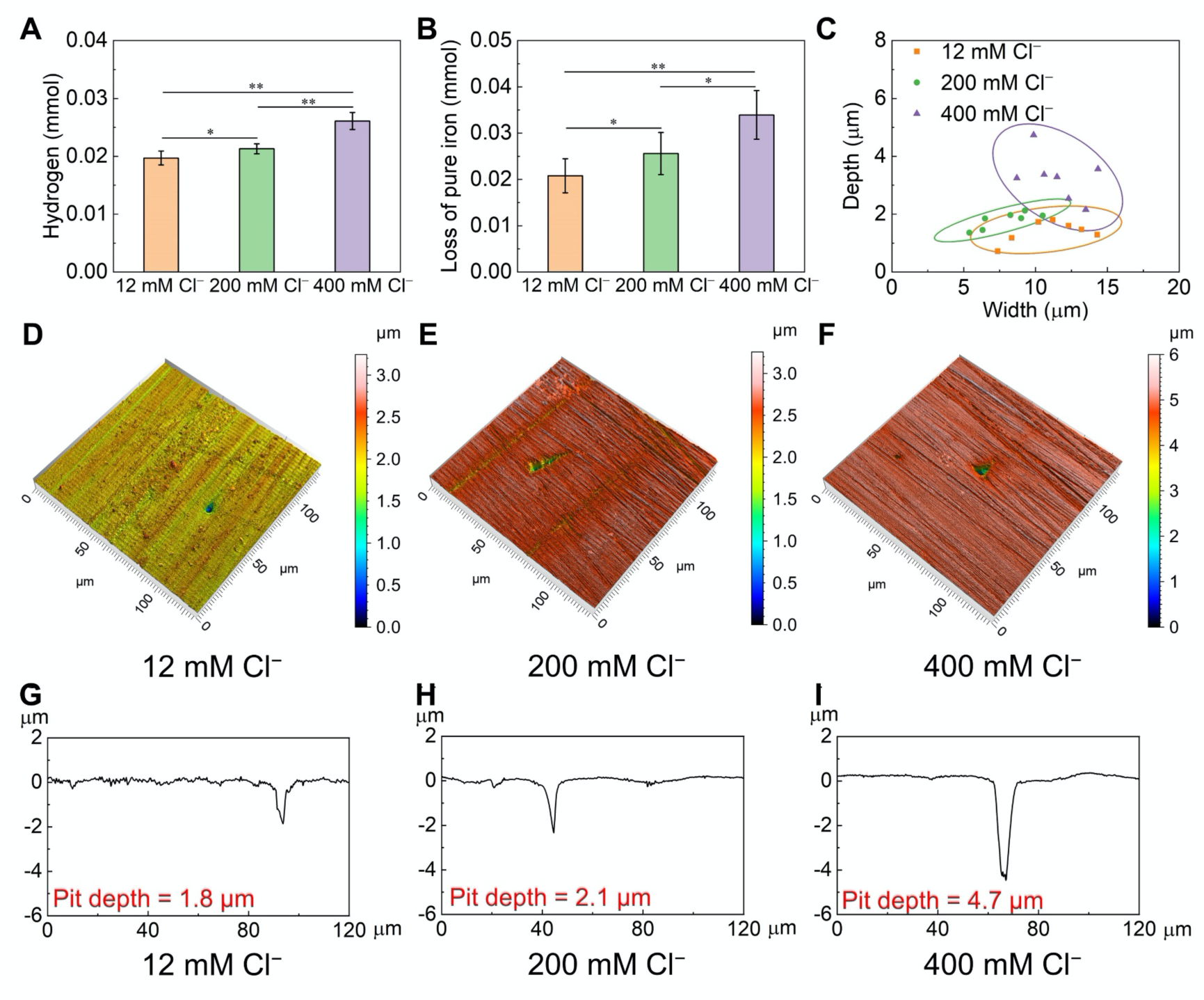
The influence of chloride concentrations on anaerobic abiotic Fe^0^ corrosion in NB culture medium (5 mM lactate and 5 mM sulfate) after seven days of incubation. (A) H_2_ accumulation (mean and standard deviation of triplicate incubations). (B) Weight loss (mean and standard deviation of triplicate incubations). (C) Depth and width of pits of the seven deepest pits in each treatment chosen from the three replicate coupons. (D-I) Surface morphology and profiles of the deepest pit for each chloride concentration. *P < 0.05, **P < 0.01.

Electrochemical parameters further confirmed that higher chloride concentrations promoted faster abiotic corrosion rates. *R*_p_, which is a measure of corrosion resistance was consistently lower over the 7 day incubation with higher chloride concentrations (Fig. 2), indicating higher corrosion rates. At day 7, corrosion current densities (*i*_corr_) were higher with higher chloride concentrations (Fig. 2, Table S1). The higher abiotic rates of corrosion with higher chloride are consistent with previously developed concepts for chloride-dependent abiotic steel corrosion [30, 37].

**Fig. 2.**
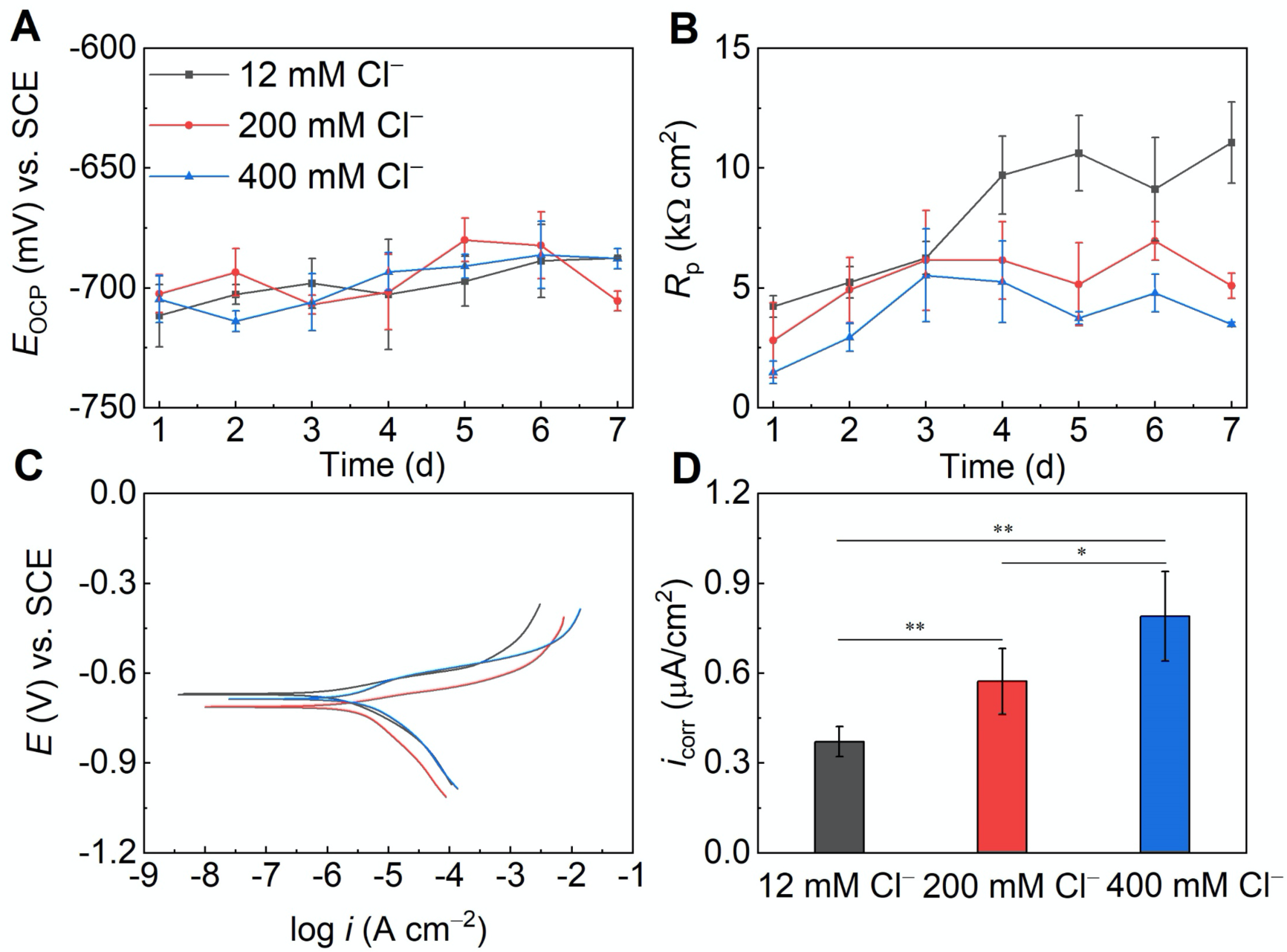
Electrochemical analysis of the influence of chloride concentrations on anaerobic abiotic Fe^0^ corrosion in NB culture medium (5 mM lactate and 5 mM sulfate). (A) Open circuit potential (OCP). (B) Polarization resistance *R*_p_ determined from linear polarization resistance measurements. (C) Potentiodynamic polarization curves after seven day immersion. (D) Corrosion current densities (*i*_corr_) calculated from (C). Mean and standard deviation of triplicate samples. *P < 0.05, **P < 0.01.

### Increased chloride accelerates Fe^0^ corrosion in the presence of *D. vulgaris*

The increase in abiotic H_2_ production associated with higher chloride concentrations is expected provide more H_2_ for sulfate reducers to reduce sulfate. As detailed in the Introduction, increased sulfide production rates associated with greater H_2_ availability is expected in turn to further enhance H_2_ production and associated Fe^0^ corrosion. Therefore, in order to determine whether increased chloride would increase corrosion rates in the presence of *D. vulgaris*, strains were adapted to grow at higher chloride concentrations. *D. vulgaris* grown in freshwater NB medium (12 mM chloride) was transferred to NB medium with a chloride concentration that was 10% of marine medium (40 mM chloride). As the culture adapted to growth at 40 mM chloride with successive transfers, it was transferred to medium with incrementally higher chloride concentrations until a strain adapted to grow at 50% of the chloride concentration of marine medium (200 mM) was obtained. This strain grew in the 200 mM chloride medium as well as the parental strain grew in the 12 mM chloride freshwater medium (Fig. 3A). With continued transfer in media with increasingly higher chloride concentrations, a strain was adapted to grow in medium with the 400 mM chloride, designated marine NB medium. This strain had a longer lag period and a slightly lower maximum cell density than the unadapted strain grown in freshwater medium, but this difference was not sufficient to substantially impact on growth in seven-day corrosion studies (Fig. 4). Biofilm formation of the three different strains at their respective chloride concentrations was similar. In each case microcolonies of cells were sparsely distributed on the Fe^0^ surface with occasional pillars (Fig. 4).

**Fig. 3.**
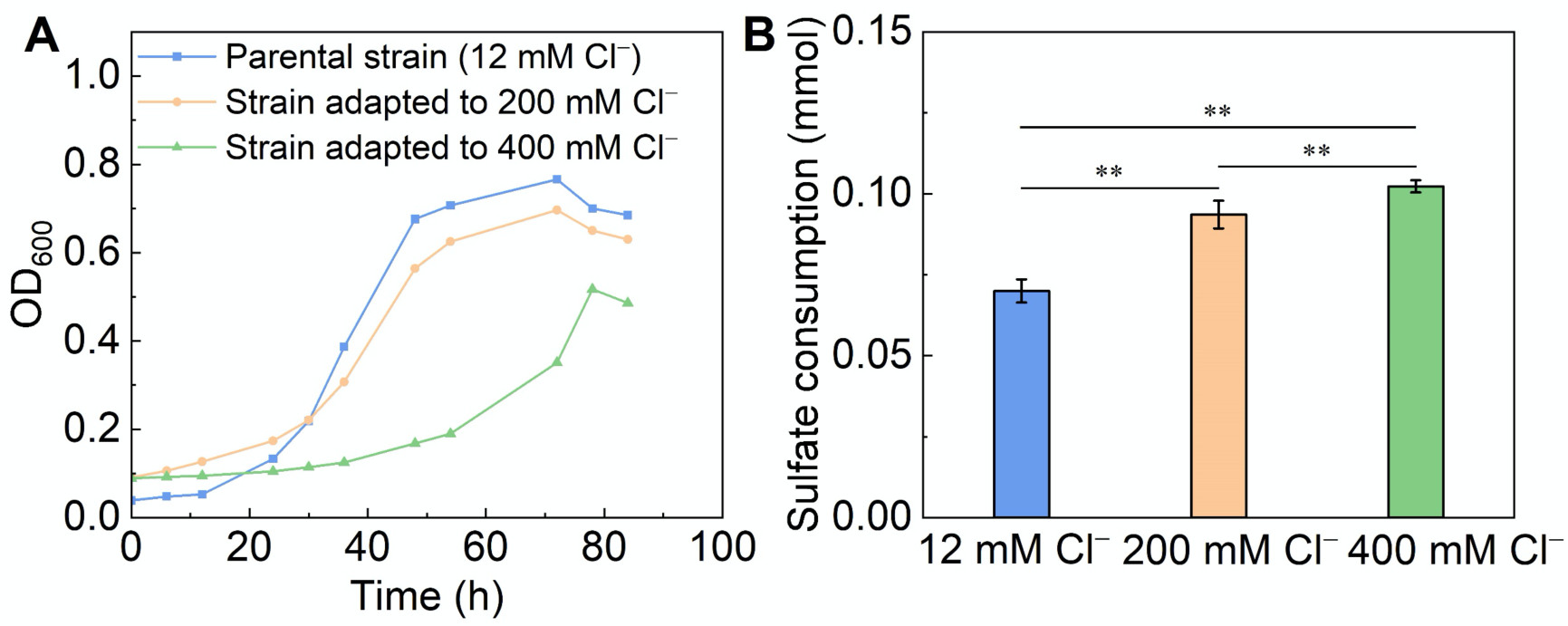
*D. vulgaris* growth and activity at different chloride concentrations. (A) Parental strain growing in freshwater medium (12 mM chloride); strain adapted to mid-level chloride growing in medium with 200 mM chloride; and strain adapted to marine chloride level growing in medium with 400 mM chloride. (B) Sulfate consumed after seven days in cultures with Fe^0^ and lactate as potential electron donors either by the parental strain in 12 mM chloride medium; strain adapted to mid-level chloride in 200 mM chloride medium; or strain adapted to marine level chloride growing in medium with 400 mM chloride. Results are the means and standard deviation of triplicate incubations. **P < 0.01.

**Fig. 4.**
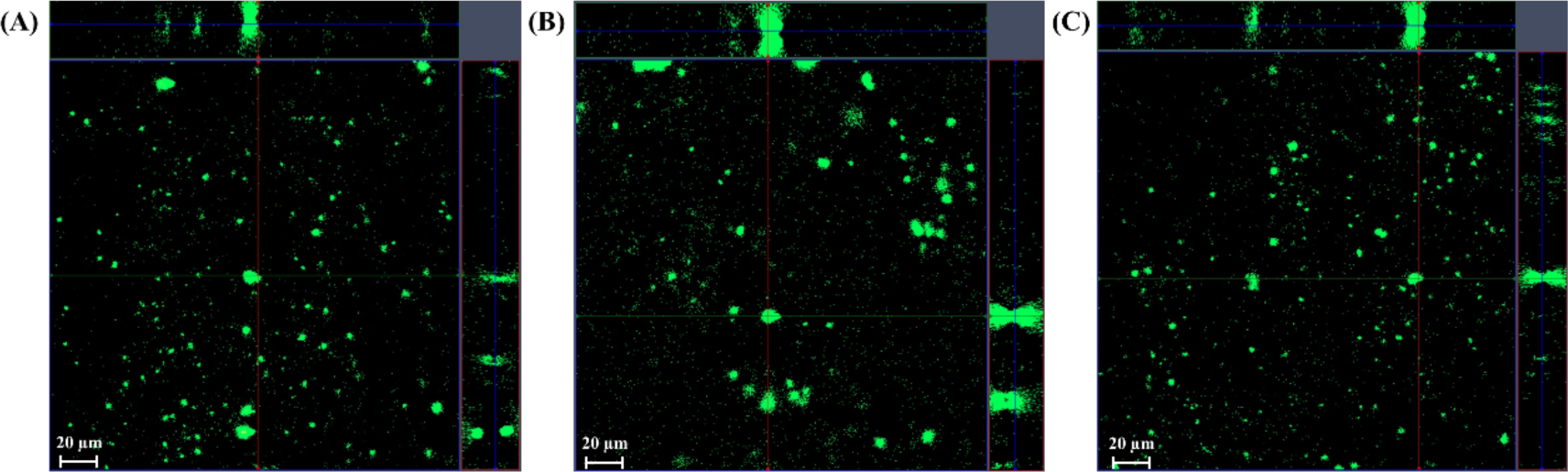
Confocal scanning laser microscopy images of *D. vulgaris* cells on Fe^0^. (A) Parental strain growing in freshwater medium (12 mM chloride). (B) Strain adapted to mid-level chloride growing in medium with 200 mM chloride. (C) strain adapted to marine chloride level growing in medium with 400 mM chloride. Large central images show x-y plane of *D. vulgaris* biofilm, small top and right side images show x-z and y-z planes of *D. vulgaris* biofilm, respectively.

*D. vulgaris* reduced more sulfate in media with higher chloride concentrations (Fig. 3B). As detailed in the Introduction, *D. vulgaris* does not directly extract electrons from Fe^0^, but rather relies on H_2_ produced from abiotic reactions [8, 14]. The most likely source of the additional electron donor to promote more sulfate reduction at higher chloride concentrations was the increased H_2_ production that the abiotic studies demonstrated were associated with more chloride. As detailed in the Introduction, any increase in H_2_ production will contribute to a positive feedback loop as the additional H_2_ produced (reaction #1) will stimulate more *D. vulgaris* sulfide production (reaction #2), which in turn will generate even more H_2_ from the positive impact of FeS on the abiotic electron transfer of Fe^0^ oxidation coupled to H_2_ production (reaction #1) as well as more direct sulfide-associated H_2_ production form Fe^0^ (reaction #3).

In accordance with this model of higher chloride promoting higher Fe^0^ corrosion in the presence of *D. vulgaris*, all physical and electrochemical measurements of the *D. vulgaris* cultures demonstrated that Fe^0^ was corroded faster at higher chloride concentrations. This data included: greater weight loss from the Fe^0^ coupons and more extensive pitting (Fig. 5), as well as a lower *R*_p_ and higher *i*_corr_ (Fig. 6, Table S2) with additional chloride. All measures of corrosion in the presence of *D. vulgaris* were much higher than in abiotic controls, demonstrating that the differences in corrosion with the different strains and chloride concentrations could be attributed in part to microbial activity, such as sulfide production, not just chloride-induced abiotic corrosion.

**Fig. 5.**
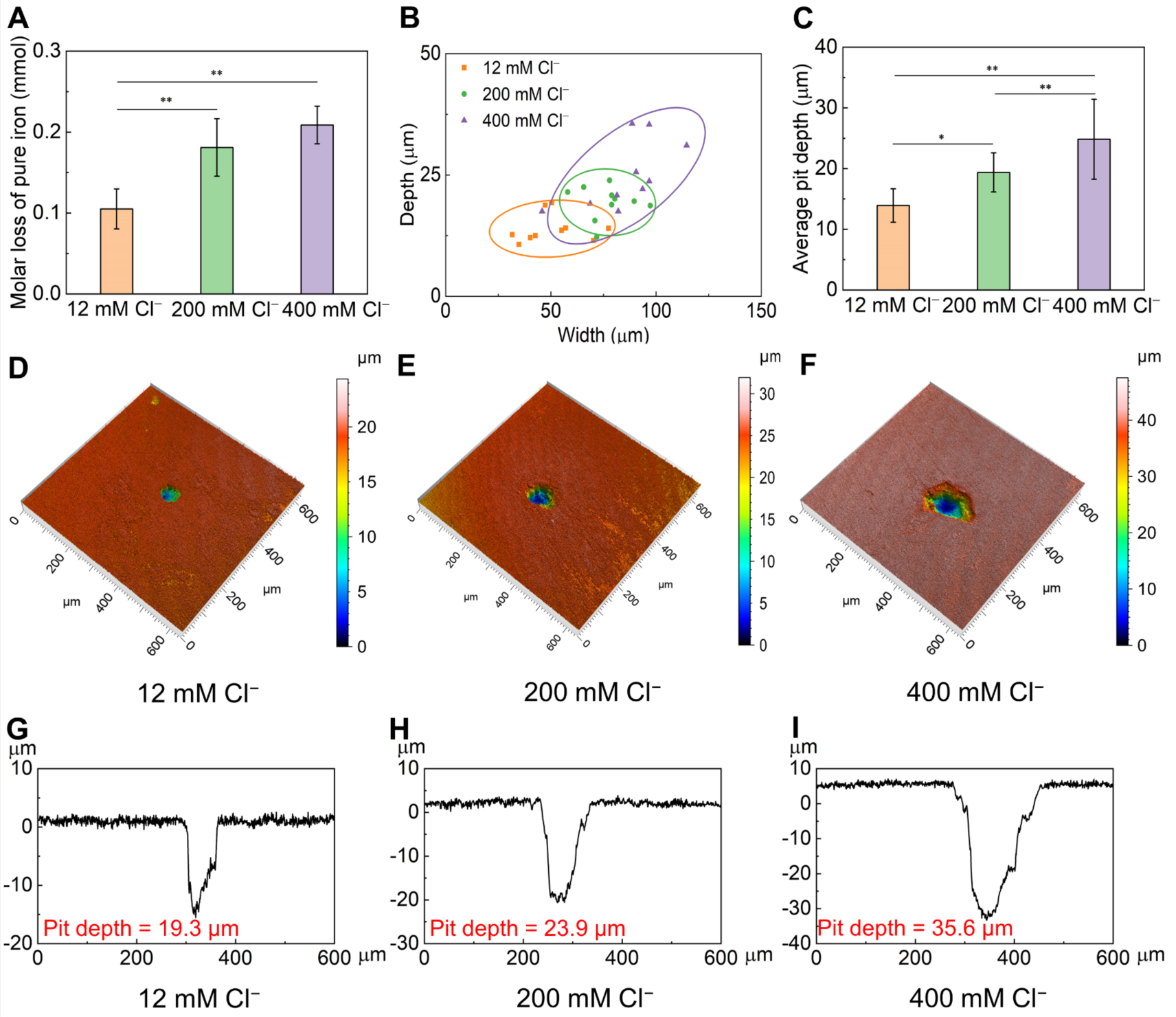
The influence of chloride concentrations on *D. vulgaris* Fe^0^ corrosion. (A) Iron loss after seven days in cultures with different chloride concentrations and strains adapted for growth at those chloride concentrations. Mean and standard deviation of triplicate cultures. (B) Depth and width of pits from the ten deepest corrosion pits on iron surfaces for each chloride concentration (C) Average pit depth. Mean and standard deviation of triplicate samples (D-I) Surface morphology and profiles of the deepest pit for each chloride concentration. *P < 0.05, **P < 0.01.

**Fig. 6.**
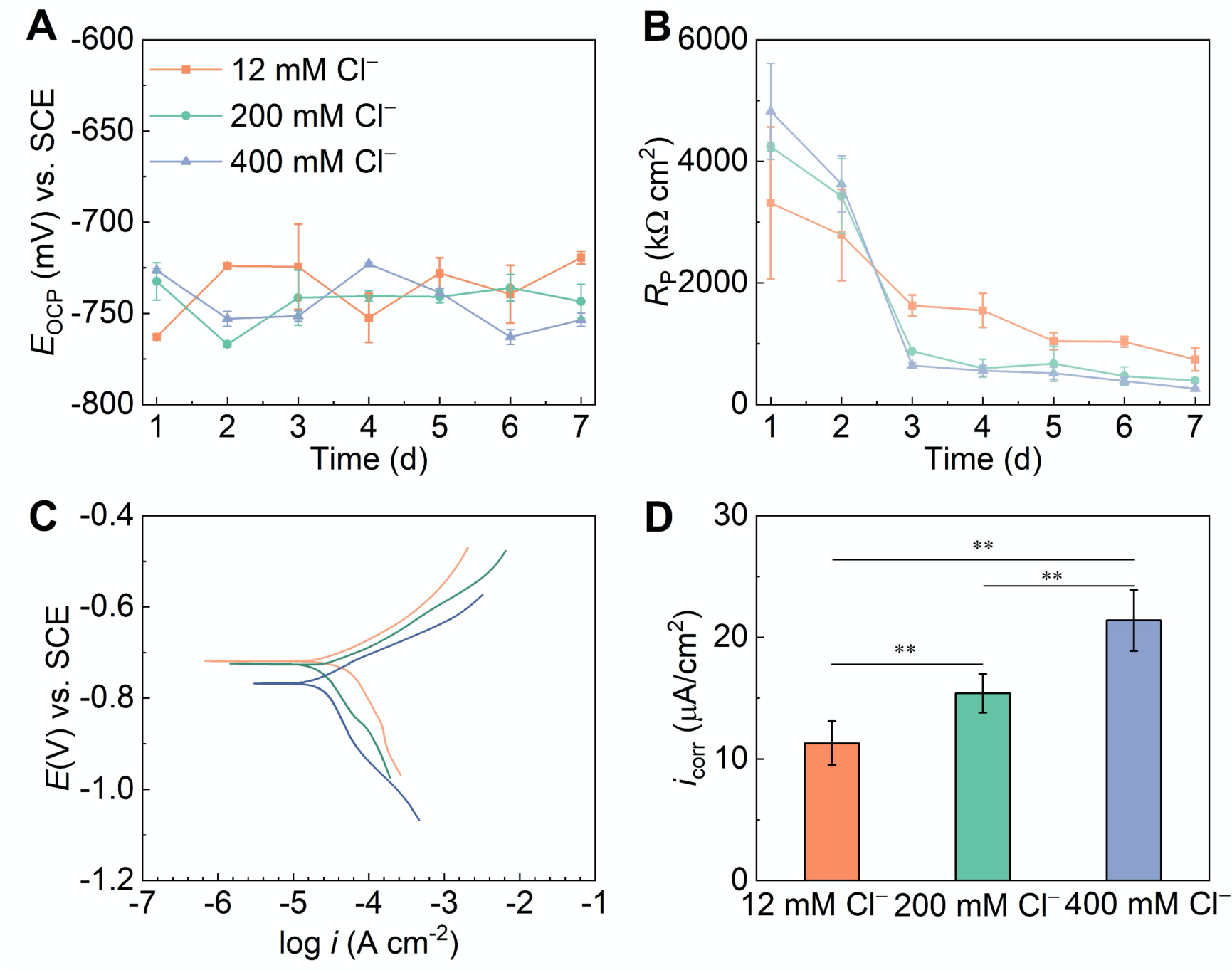
Electrochemical analysis of the influence of chloride concentrations on *D. vulgaris* Fe^0^ corrosion in NB culture medium (5 mM lactate and 5 mM sulfate). (A) Open circuit potential (OCP). (B) Polarization resistance *R*_p_ determined from linear polarization resistance measurements by LPR measurement during 7-d incubation. (C) Potentiodynamic polarization curves after seven days of incubation. (D) Corrosion current densities (*i*_corr_) calculated from (C). Mean and standard deviation of triplicate samples. **P < 0.01.

Analysis of corrosion products with energy dispersive X-ray spectroscopy revealed small differences in elemental distribution, but X-ray diffraction found no differences in the abundance of iron sulfide or iron oxide corrosion products (Fig. 7). Thus, chloride’s impact on corrosion rates did not appear to be related to differences in corrosion product formation.

**Fig. 7.**
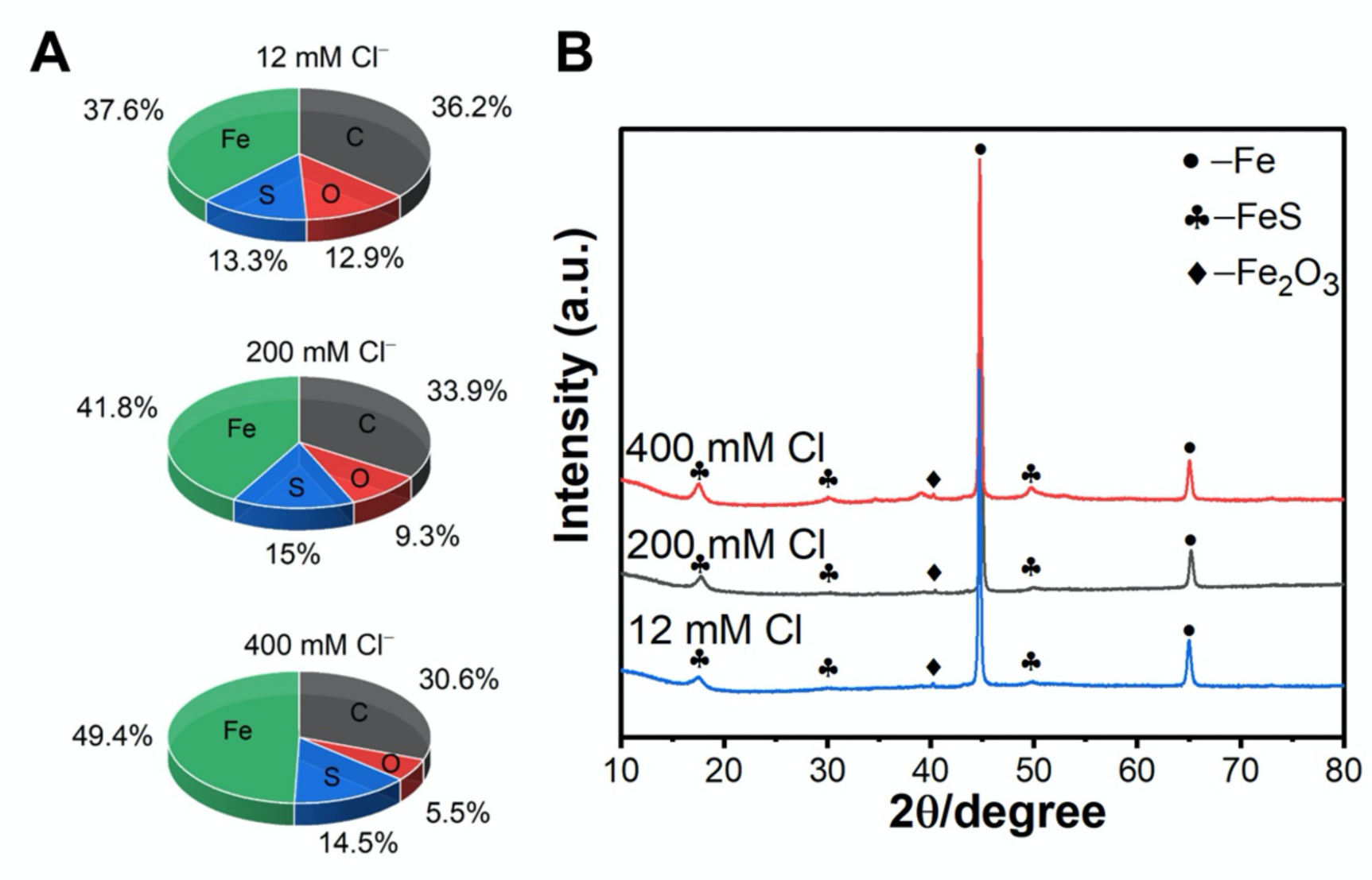
Elemental composition and corrosion products on iron surfaces. (A) Energy-dispersive X-ray spectroscopy elemental analysis. (B) X-ray diffraction analysis of corrosion products.

Corrosion rates in the presence of *D. ferrophilus* are media-dependent.

As detailed in the Introduction, *D. ferrophilus* has been reported to corrode iron-containing metals faster than *D. vulgaris*. To determine whether these differences in corrosion rates could be attributed to differences in medium composition, we attempted to adapt *D. vulgaris* to grow in the 195C high chloride medium routinely used to culture *D. ferrophilus*, but were unsuccessful. *D. ferrophilus* was adapted to grow in the 400 mM chloride marine NB medium employed in the *D. vulgaris* studies with lactate as the electron donor and sulfate as the electron acceptor (Fig. 8A), growing better than *D. vulgaris* in this same marine medium (Fig. 3A). *D. ferrophilus* effectively colonized Fe^0^ when it was included the marine NB medium (Fig. 8B). However, Fe^0^ weight loss (Fig. 8C), pitting size (Fig. 8D, E) and maximum pit depths (Fig. 5F, I versus Fig. 8F, G) were greater in *D. vulgaris* cultures. Electrochemical analysis also indicated that Fe^0^ was corroded faster in the presence of *D. vulgaris* than *D. ferrrophilus* in the same marine NB medium (Fig. 9, Table S3).

**Fig. 8.**
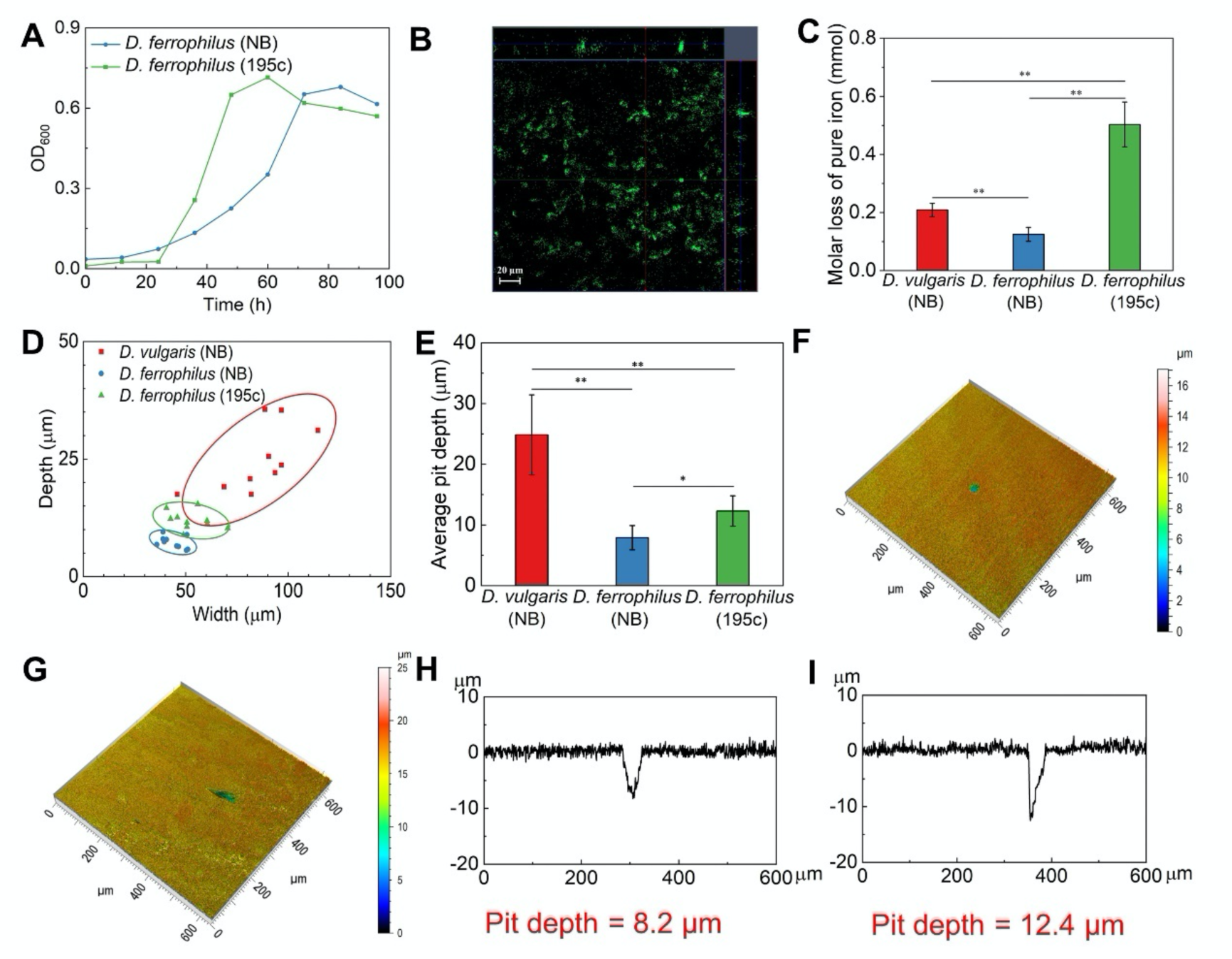
*D. ferrophilus* growth and corrosion in either marine NB medium (400 mM chloride) or 195c medium (400 mM chloride) compared with *D. vulgaris* grown in marine NB medium. (A) Comparison of *D. ferrophilus* growth in marine NB and 195C media. (B) Confocal micrograph of *D. ferrophilus* biofilm on Fe^0^ after seven days growth in marine NB medium. Large central image shows the x-y plane of *D. ferrophilus* biofilm, small top and side images show x-z plane and y-z plane of *D. ferrophilus* biofilm, respectively. (C) Fe^0^ weight loss after seven days for *D. vulgaris* in marine NB medium (from Fig. 5A) and *D. ferrophilus* in marine NB and 195C media. (Mean and standard deviation of triplicate samples.) (D) Depth and width of pits from the ten deepest corrosion pits on iron surfaces for *D. vulgaris* in NB medium (from Fig. 5B) and *D. ferrophilus* in marine NB and 195C medium. (E) Average pit depth of *D. vulgaris* in NB medium (from Fig. 5C) (Mean and standard deviation of triplicate samples.) and *D. ferrophilus* in marine NB and 195C medium. (F-I) maximum pit depth for *D. ferrophilus D. ferrophilus* in marine NB and 195C medium. *P < 0.05, **P < 0.01.

**Fig. 9.**
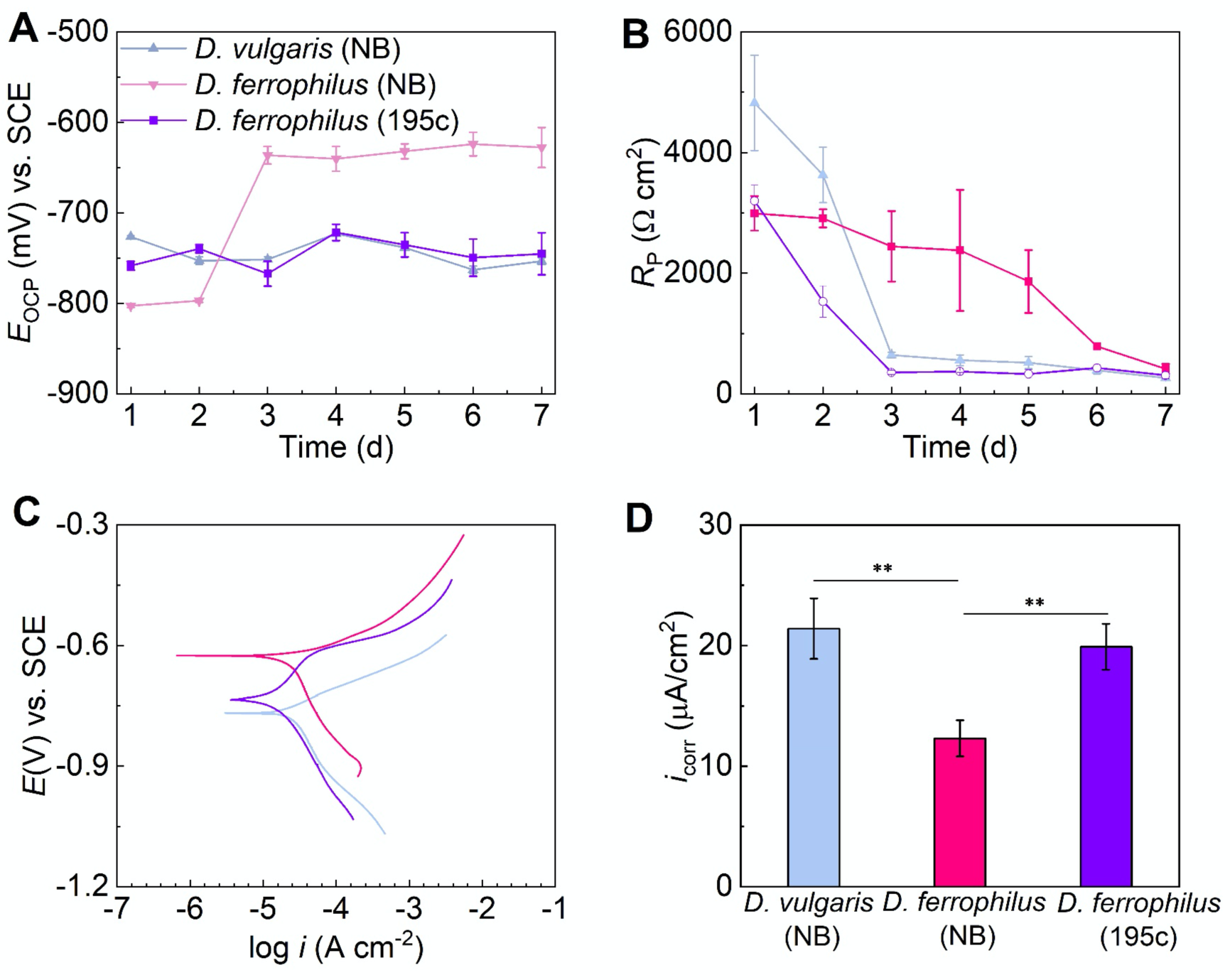
Electrochemical analysis of *D. ferrophilus* corrosion grown in either marine NB medium (400 mM chloride) or 195c medium (400 mM chloride) compared with *D. vulgaris* grown in marine NB medium. (A) Open circuit potential (OCP). (B) Polarization resistance *R*_p_ determined from linear polarization resistance measurements by LPR measurement during 7-d incubation. (C) Potentiodynamic polarization curves after seven days of incubation. (D) Corrosion current densities (*i*_corr_) calculated from (C). Mean and standard deviation of triplicate samples. **P < 0.01.

Corrosion in the presence of *D. ferrophilus* was also evaluated in the 195C medium used in previous corrosion studies [23, 38]. This medium differs slightly from NB medium in the composition of major ions and trace metals, but has the same chloride concentration as the ‘marine’ NB medium. The weight loss when *D. ferrophilus* was cultured in 195C medium was more than twice the weight loss when *D. vulgaris* was grown in marine NB medium (Fig. 8C), but pitting was substantially greater in the *D. vulgaris* cultures (Fig.5F, I, Fig. 8D, E, F, I). Corrosion current associated with *D. vulgaris* growing in NB medium were slightly higher than corrosion current for *D. ferrophilus* grown in 195C medium. These results indicate that slight differences in media composition can impact on corrosion rates in the presence of *D. ferrophilus*, but that corrosion is not greater when *D. ferrophilus* and *D. vulgaris* are grown at high chloride concentrations in the same medium.

## Conclusion

The results demonstrate that Fe^0^ corrosion rates in the presence of sulfate-reducing bacteria are highly dependent upon the medium composition. Chloride concentration was a major factor in the corrosion rates in *D. vulgaris* cultures. Further study is required to evaluate the potential impact of other medium constituents on corrosion. For example, inorganic carbon and/or phosphate are typically added in high concentrations to buffer anaerobic growth medium. Both may have either positive or negative impacts on corrosion rates by influencing the formation of corrosion products and contributing to the formation of passivation layers and/or influencing microbial growth. Even medium constituents added at low concentrations, such as vitamins and trace metals, might influence corrosion rates by enhancing the synthesis of enzymes or co-factors and/or acting as electron shuttles to promote electron transfer between Fe^0^ and cells. Thus, careful consideration of medium composition is required in any studies attempting to characterize the corrosion rates associated with different microbes.

The initial *D. ferrophilus* corrosion studies compared *D. ferrophilus* grown in marine medium with *D. vulgaris* grown in freshwater medium [21]. The results presented here demonstrate that differences in chloride concentrations were likely a major factor in the faster corrosion in *D. ferrophilus* cultures reported in those studies, which were mistakenly attributed to *D. ferrophilus* direct electron uptake from Fe^0^. Even when medium compositions can be matched, comparing corrosion rates between microbes is not a valid approach for elucidating differences in corrosion mechanisms for several reasons [1, 5, 24, 39]. Functional genetic approaches with the appropriate mutant strains are required for rigorous evaluation of corrosion mechanisms [1, 5, 14].

The composition of typical cultivation medium is substantially different from the concentrations of electron donors and nutrients that microbes experience in typical environments in which metal corrosion is prevalent. These considerations suggest that future laboratory microbial corrosion studies, including functional genetic studies, should attempt to mimic environmental conditions as much as possible in order to obtain environmentally relevant results.

## Acknowledgement

This work was financially supported by the National Key Research and Development Program of China (No. 2022YFB3808800), China Baowu Low Carbon Metallurgy Innovation Foundation-BWLCF202120, and the National Natural Science Foundation of China (No. 52301080).

## Notes

### Competing Interest Statement

The authors have declared no competing interest.

